# CoV3D: A database and resource for high resolution coronavirus protein structures

**DOI:** 10.1101/2020.05.12.091983

**Authors:** Ragul Gowthaman, Johnathan D. Guest, Rui Yin, Jared Adolf-Bryfogle, William R. Schief, Brian G. Pierce

## Abstract

SARS-CoV-2, the etiologic agent behind COVID-19, exemplifies the general threat to global health posed by coronaviruses. The urgent need for effective vaccines and therapies is leading to a rapid rise in the number of high resolution structures of SARS-CoV-2 proteins that collectively reveal a map of virus vulnerabilities. To assist structure-based design of vaccines and therapeutics against SARS-CoV-2 and other coronaviruses, we have developed CoV3D, a database and resource for coronavirus protein structures, which is updated on a weekly basis. CoV3D provides users with comprehensive sets of structures of coronavirus proteins and their complexes with antibodies, receptors, and small molecules. Integrated molecular viewers allow users to visualize structures of the spike glycoprotein, which is the major target of neutralizing antibodies and vaccine design efforts, as well as sets of spike-antibody complexes, spike sequence variability, and known polymorphisms. In order to aid structure-based design and analysis of the spike glycoprotein, CoV3D permits visualization and download of spike structures with modeled N-glycosylation at known glycan sites, and contains structure-based classification of spike conformations, generated by unsupervised clustering. CoV3D can serve the research community as a centralized reference and resource for spike and other coronavirus protein structures, and is available at: https://cov3d.ibbr.umd.edu.

## Introduction

Coronaviruses (CoVs) have been responsible for several outbreaks over the past two decades, including SARS-CoV in 2002-2003, MERS-CoV in 2012 (1), and the current COVID-19 pandemic, caused by SARS-CoV-2, which began in late 2019 (2). The scale of the COVID-19 pandemic has led to unprecedented efforts by the research community to rapidly identify and test therapeutics and vaccines, and to understand the molecular basis of SARS-CoV-2 entry, pathogenesis, and immune targeting.

Since February 2020, a large number of SARS-CoV-2 protein structures have been released in the Protein Data Bank (PDB) (3). As of June 17th, 2020, this includes 28 spike glycoprotein structures, over 150 main protease structures, and over 60 structures of other SARS-CoV-2 proteins. These high-resolution protein structures are of immense importance for understanding viral assembly and to aid rational vaccine and therapeutic design. The first structures of the SARS-CoV-2 trimeric spike glycoproteins (the major target of SARS-CoV-2 vaccines and antibody therapeutics) were reported in February and early March 2020 (4,5). Previously determined spike glycoprotein structures have enabled advances including rational stability optimization of SARS-CoV and MERS-CoV spikes, yielding improved protein expression and immunogenicity (6). Given that the rapid rate of coronavirus protein structural determination and deposition is likely to continue, a simple and updated resource detailing these structures would provide a useful reference.

Here we describe a new database of experimentally determined coronavirus protein structures, CoV3D. CoV3D is updated automatically on a weekly basis, as new structures are released in the PDB. Structures are classified by CoV protein, as well as bound molecule, such as monoclonal antibody, receptor, and small molecule ligand. To enable insights into the spike glycoprotein, we also include information on SARS-CoV-2 residue polymorphisms, overall coronavirus sequence diversity of betacoronaviruses mapped onto spike glycoprotein structures, and structures of spike glycoproteins with modeled glycans, as a reference or for subsequent modeling. This resource can aid in efforts for rational vaccine design, targeting by immunotherapies, biologics, and small molecules, and basic research into coronavirus structure and recognition. CoV3D is publicly available at https://cov3d.ibbr.umd.edu.

## Methods

### Web and database implementation

CoV3D is implemented using the Flask web framework (https://flask.palletsprojects.com/) and the SQLite database engine (https://www.sqlite.org/).

### Structure identification, visualization, and glycan modeling

Structures are identified from the PDB on a weekly basis using NCBI BLAST command line tools (7), with coronavirus protein reference sequences from SARS-CoV, MERS-CoV, and SARS-CoV-2 as queries. The spike glycoprotein reference sequences (GenBank identification NP_828851.1, YP_009047204.1 and QHD43416.1 for SARS-CoV, MERS-CoV and SARS-CoV-2 virus respectively) are used as queries to identify all available spike glycoprotein structures. Peptide-MHC structures containing coronavirus peptides are identified in the PDB through semi-manual searches of the PDB site and literature, though future automated updates are planned in conjunction with an expanded version of the TCR3d database (8). Structural visualization is performed using NGL viewer (9). N-glycans are modeled onto spike glycoprotein structures using a glycan modeling and refinement protocol in Rosetta (10). An example command line and Rosetta Script for this glycan modeling protocol is provided as Supplemental Information.

### Spike clustering and classification

Root-mean-square distances (RMSDs) between all pairs of full CoV spike glycoprotein chains were computed using the FAST structure alignment program (11). The resultant distance matrix was input to R (www.r-project.org) which was used to perform hierarchical clustering, and the dendrogram was generated using the dendextend R package (12). The spike chains were classified into two clusters based on this analysis, corresponding to open and closed spike states.

### Sequence data collection and analysis

SARS-CoV-2 spike glycoprotein sequences were downloaded from NCBI Virus (13), followed by filtering out sequences with missing residues. Sequence polymorphism information was obtained by BLAST search using a reference SARS-CoV-2 spike glycoprotein sequence (QHD43416.1). To develop spike glycoprotein alignments, betacoronavirus spike glycoprotein sequences were downloaded from NCBI Virus (13) and aligned with Clustal Omega (14) in SeaView (15). Sequences that were redundant (>95% similarity) or contained missing residues were removed, with the remaining 70 sequences forming the Pan-betacoronavirus alignment. A subset of 18 sequences from the pan-betacoronavirus alignment was used to generate the SARS-like sequence alignment, which contains every sequence from the pan-betacoronavirus alignment with >70% sequence similarity to the SARS-CoV-2 spike. Sequence logos are generated dynamically for user-specified residue ranges using the command-line version of WebLogo (16). Phylogenetic trees representing spike reference sequences were generated using ClustalX (17), and visualized using the APE package (18) in R (www.r-project.org).

## Results

### Database contents

The main components of the CoV3D database are interrelated tables, datasets, and tools for coronavirus protein structures and spike glycoprotein sequences. A schematic of the CoV3D input, organization, and contents is shown in **Figure 1**. The structure portion of the database includes dedicated pages and tables for:

- Spike glycoprotein structures
- Spike conformational classification
- Antibody structures
- Spike structures with modeled glycans
- Main protease structures
- Other SARS-CoV-2 protein structures (nucleocapsid, NSPs, ORFs, etc.)
- Peptide-Major Histocompatibility Complex (MHC) structures with coronavirus peptides

**Figure 1.**
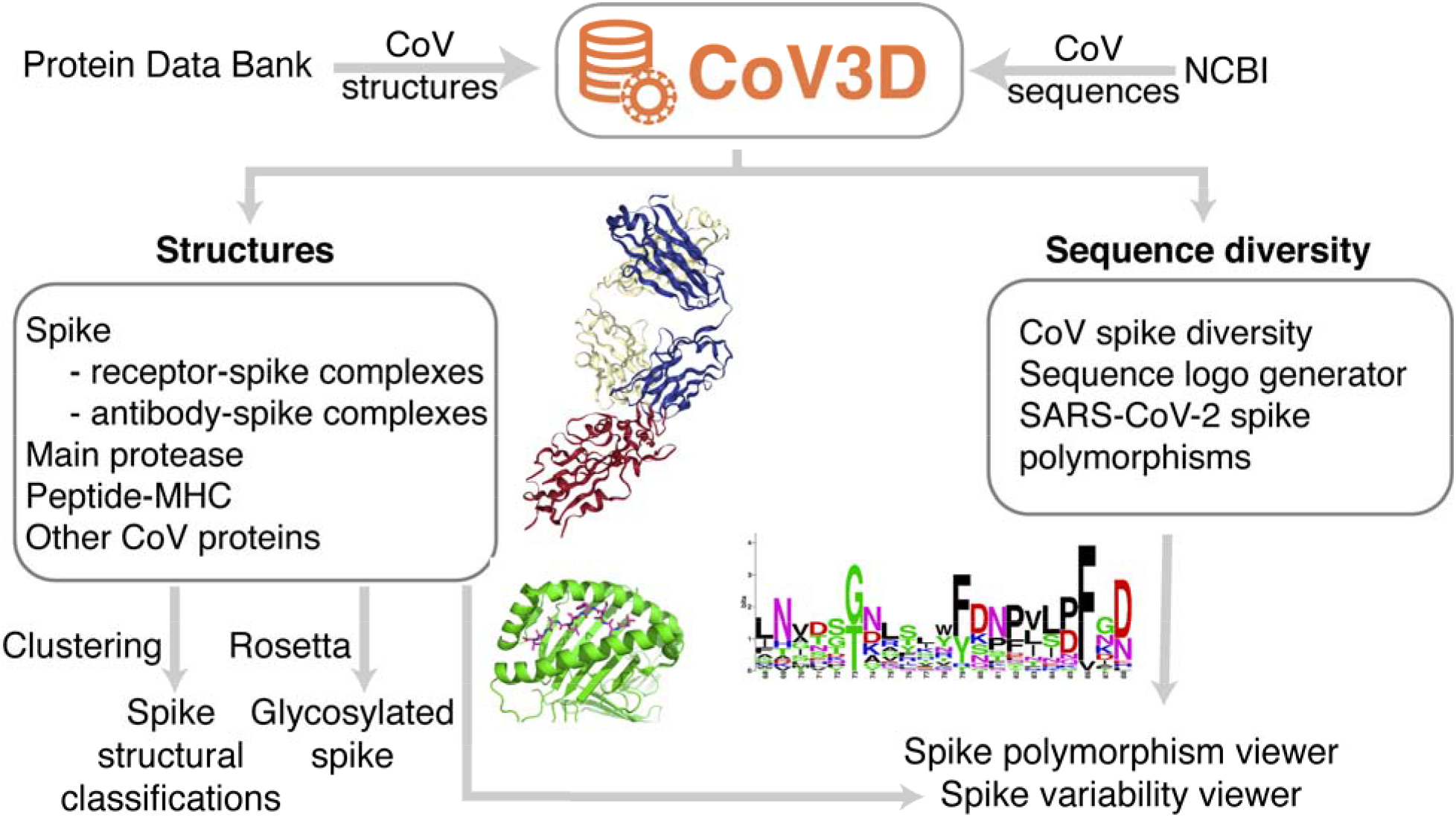
Implementation and organization of CoV3D. The database combines curated and annotated CoV structural data from the PDB, updated automatically on a weekly basis, as well as CoV sequence data from NCBI. Structures include immunologically relevant complexes; shown are an antibody in complex with the SARS-CoV-2 spike receptor binding domain (PDB code 7BZ5) (25), and a SARS-CoV membrane protein peptide in complex with the human major histocompatibility complex (MHC) protein HLA-A2 (PDB code 3I6K) (26). Also shown is a sequence logo generated in CoV3D for a spike glycoprotein subsequence.

Spike glycoprotein structures are annotated by binding partner(s), including bound antibody and receptor, as well as domains present in the structure. Protease structures include annotation of bound ligand, for convenience of those investigating protease inhibitors computationally or experimentally. Antibody structures are annotated by complementarity determining region (CDR) loop clusters, assigned by PyIgClassify (19).

Spike conformational classifications are performed for each chain in structures containing full spike glycoproteins with a receptor binding domain (RBD) present in the structure. Unsupervised clustering based on spike root-mean-square distances (RMSDs) automatically classifies the spike structures into two broad clusters (**Figure 2**), corresponding to open and closed conformations (also referred to as “up” and “down” conformations); these forms are relevant for receptor binding and vaccine design (4,5), and annotations of those states based on this analysis are included in CoV3D. Out of 129 SARS-CoV-2, SARS-CoV, and MERS-CoV spike structures in the current set, 39 of them are assigned to the open state, while the remaining 90 are closed (**Figure 2**).

**Figure 2.**
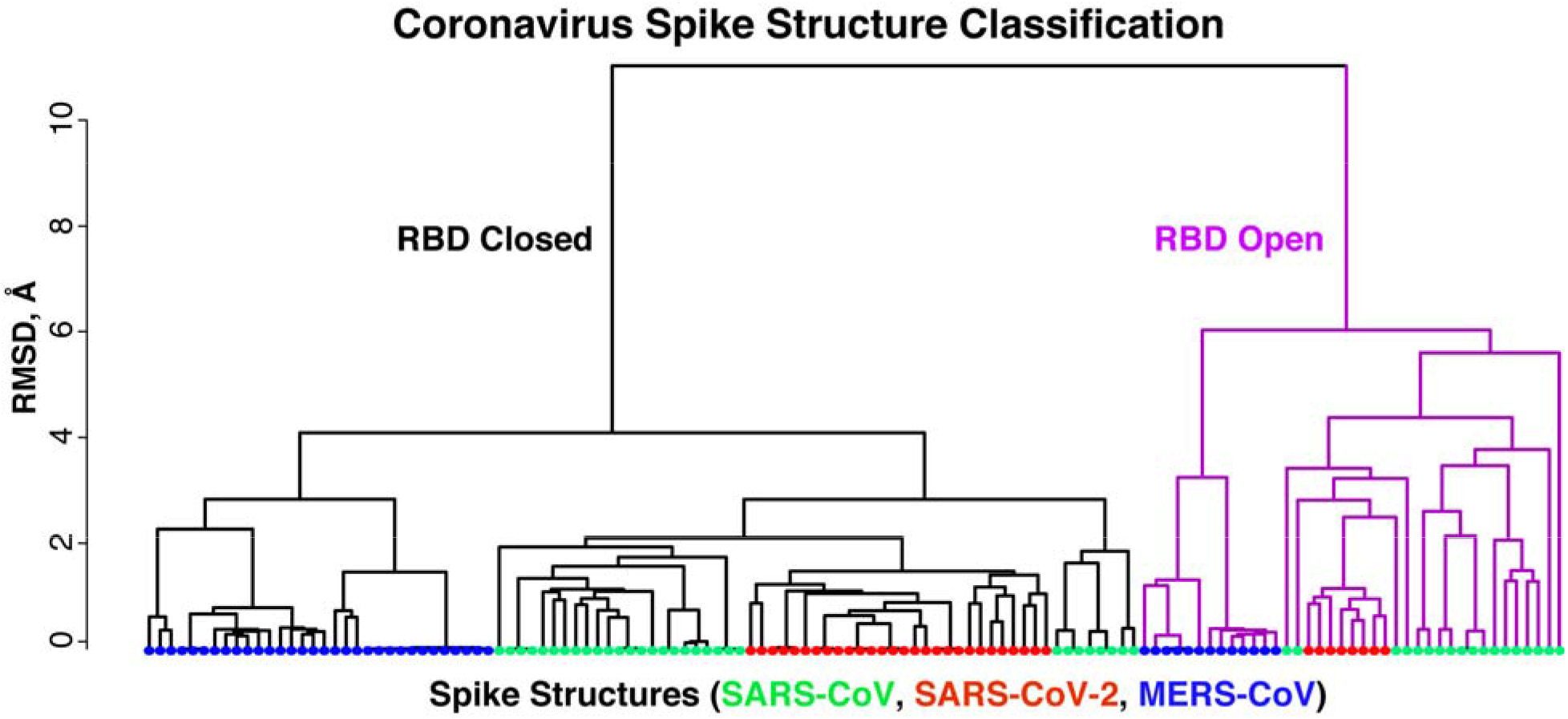
Structural clustering of coronavirus spike glycoprotein structures. The dendrogram of was generated based on pairwise structural similarities between 129 spike glycoprotein chain structures in R (www.r-project.org). Leaves are colored by virus of the spike glycoprotein structure (SARS-CoV, SARS-CoV-2, or MERS-CoV), and branches are colored magenta for the cluster corresponding to receptor binding domain (RBD) open spike conformations. This clustering is used to assign open and closed states to spike structures in CoV3D.

The sequence section of CoV3D focuses is focused on spike glycoprotein variability, and includes:

- Spike glycoprotein residue polymorphisms
- Spike glycoprotein sequence logos based on sets of betacoronavirus sequences
- Visualization of conservation on spike glycoprotein structures

Additional features of the database include a page with recently released structures, and a downloads page for users to download structural data, sequence data, and tables. Summary statistics are also given, providing metrics such as number of spike structures released for SARS-CoV, MERS-CoV, and SARS-CoV-2.

### Example use case: viewing sarbecovirus conservation in an RBD antibody epitope

In addition to a table of all known structures of CoV spike proteins and their interactions, CoV3D provides a viewer with SARS-CoV-2 spike RBD interactions with antibodies and the ACE2 receptor superposed by RBD in a common reference frame (**Figure 3A**). This viewer permits users to assess overlapping or shared RBD binding modes among antibodies and ACE2. One antibody of interest is S309, which is a human antibody that was cloned from an individual who was infected with SARS-CoV in 2003, and potently neutralizes SARS-CoV-2 (20). Comparing its RBD binding with ACE2 in the RBD interaction viewer (**Figure 3A**) highlights how S309 engages the RBD at a distinct site from ACE2, in accordance with their lack of observed RBD binding competition (20). The epitope targeted by S309 is dominated by RBD residues 334-346 (circled in **Figure 3A**). On the CoV3D sequence logo generator page, users can enter spike residue ranges, and generate sequence logos that represent positional residue propensities and conservation for regions of interest, based on sets of CoV spike sequences. A sequence logo generated from a reference set of spike proteins from 18 SARS-like CoVs is shown in **Figure 3B**, indicating the conservation of the N-glycosylation site (NxS/NxT sequon) at position 343, and amino acid variability at position 340, which is glutamic acid (E) in SARS-CoV-2. Both the N343 glycan and the E340 side chain are directly engaged by S309 in the recently determined complex structure (20).

**Figure 3.**
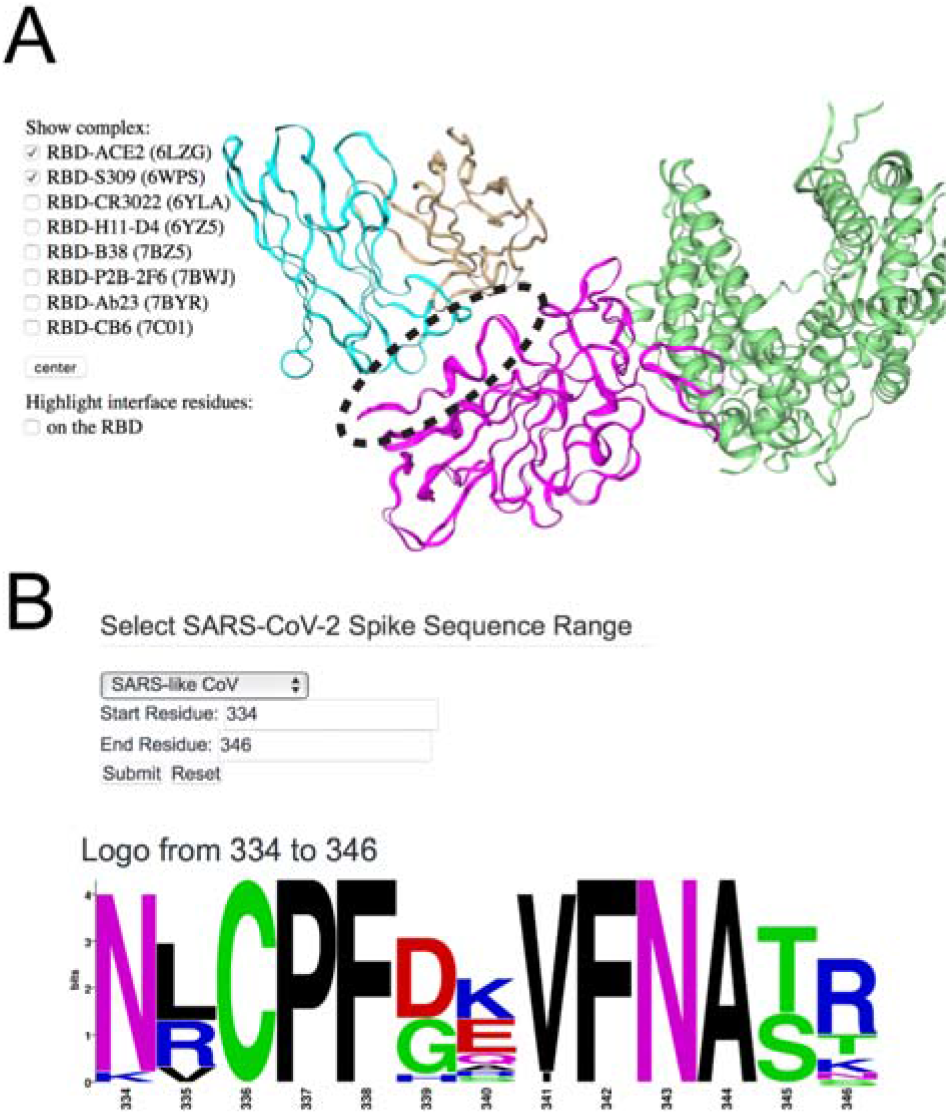
Viewing the structure, targeting, and sequence diversity of SARS-CoV-2 spike receptor binding domain (RBD). (A) Visualization of the superposed spike RBD complexes with antibody S309 (PDB code 6WPS) (20) and ACE2 receptor (PDB code 6LZG) (27) in CoV3D, with spike colored magenta, S309 heavy and light chains in cyan and tan, respectively, and ACE2 green. An interacting region of interest on the RBD (residues 334-346) is circled for reference. For clarity, only the S309-RBD region of the S309-spike complex structure is shown in the viewer. (B) CoV3D sequence logo generation interface, and logo for spike residues 334-346. The logo was generated using 18 SARS-related spike glycoprotein reference sequences and the command-line version of WebLogo (16).

### Example use case: visualizing glycosylation

N-glycosylation of viral glycoproteins can play a key role by masking the glycoprotein from the immune system, or affecting function (21). Experimentally reported protein structures often lack N-glycans, or large portions thereof, due to limitations from resolution or intrinsic glycan dynamics or heterogeneity. To enable visualization and additional analysis or modeling of glycosylated spike glycoproteins, CoV3D includes sets of structures with modeled N-glycans at all predicted glycosylation sites, with N-glycans built onto the glycoprotein structures and refined using new tools in Rosetta based on previously developed glycan functionality (10). Examples of glycosylated structures that can be visualized in CoV3D are shown in **Figure 4**, and these can be downloaded directly by users for further processing. This permits users to view features such as the N-glycosylation present on the ACE2 surface (**Figure 4A**) and the relatively high glycosylation of the spike glycoprotein stem (bottom in **Figure 4B**). Presently, these structures include uniform oligomannose glycoforms that were found to be prevalent on the SARS-CoV spike based on previous mass spectroscopy analysis (22), with five branched mannose sugars. In the future, we plan to include more spike structures with modeled glycans, and to possibly include glycoforms that reflect mass spectroscopy experimental characterization of the SARS-CoV-2 spike (23).

**Figure 4.**
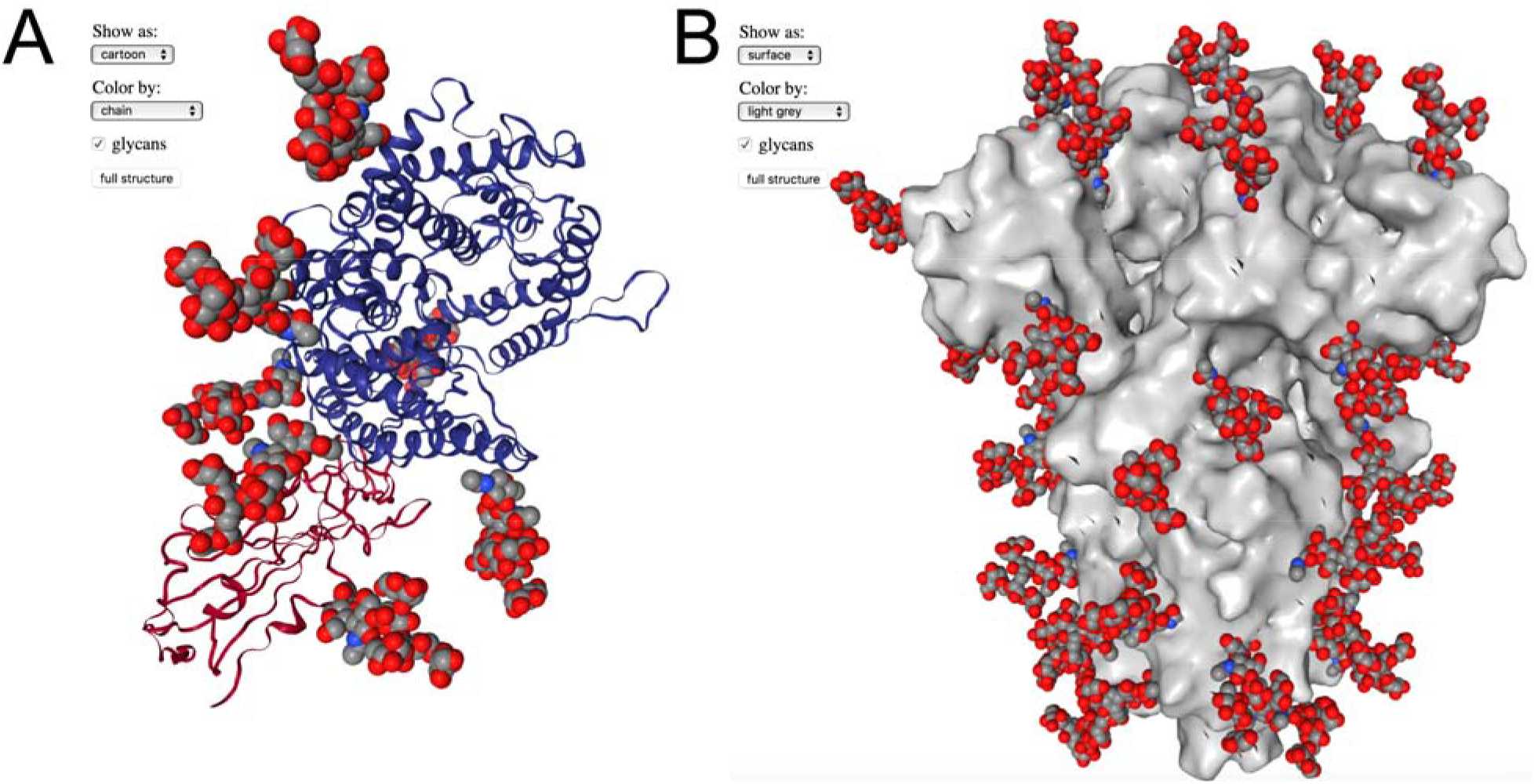
Spike structures with modeled oligomannose N-glycans. (A) The SARS-CoV-2 spike RBD-ACE2 complex (27) (PDB code 6LZG) with RBD and ACE2 shown as red and blue cartoons, respectively, with modeled N-glycans shown as gray, red, and blue spheres. One N-glycan is modeled on the spike RBD and six are modeled on ACE2. (B) A trimeric spike structure in RBD-closed conformation (5) (PDB code 6VXX) with spike shown as gray surface and modeled N-glycans shown as gray, red, and blue spheres. 48 modeled N-glycans are present on the spike structure.

## Discussion

We have constructed the CoV3D database as a reference for the research community, providing a simple and updated interface to all coronavirus 3D structures, with integrated molecular viewers, structural classification, and a variety of other useful features. This will allow researchers to identify new coronavirus protein structures as they are released, particularly for SARS-CoV-2, while enabling insights into spike glycoprotein antibody recognition, sequence features, polymorphisms, and glycosylation. Recently released SARS-CoV-2 spike-antibody complex structures have revealed mechanisms of cross-reactive CoV spike targeting and neutralization (20,24,25), and many ongoing antibody identification and characterization studies are likely to result in many more antibody-spike structures in the near future. CoV3D facilitates the use of these growing structural datasets in analysis, modeling, and structure-based design efforts by collecting and annotating these structures and their conformations, and by providing inline structural viewing of single and multiple complexes.

## Acknowledgements

We are grateful to the structural biologists and researchers whose work resulted in the structures enabling the development of this database. We are additionally thankful to Ghazaleh Taherzadeh, Stefan Ivanov, and John Moult (University of Maryland Institute for Bioscience and Biotechnology Research), as well as the RosettaCommons community, for helpful discussions and suggestions. The Institute for Bioscience and Biotechnology Research computing facility and staff, including Christian Presley, provided resources and assistance with server implementation and web hosting. This work was supported in part by NIH R01 GM126299 (to BGP).

