## Supplemental Methods for "CoV3D: A database and resource for high resolution coronavirus protein structures"

**Supplementary Methods**

**Modeling of N-glycans on spike glycoprotein structures**

Rosetta version 3.12 was used to add N-glycans to glycoprotein structures. Prior to processing with Rosetta, N-glycans present in the input structure were removed. A sample Rosetta command line for glycan addition and minimization follows:

~/rosetta/main/source/bin/rosetta_scripts.linuxgccrelease -database ~/rosetta/main/database @flags -s 6VSB.clean.pdb

Contents of the "flags" file with command line flags for the Rosetta executable are as follows:

```
-parser:protocol glycos.xml
-beta
-include_sugars
-write_pdb_link_records
-write_glycan_pdb_codes
-output_alternate_atomids
-ex1
-ex2
-overwrite
-renumber_pdb F
```

The contents of the RosettaScript file "glycos.xml" detailing the glycan structures, glycosylation sites, and protocol are as follows:

```
<ROSETTASCRIPTS>
        <SCOREFXNS>
        </SCOREFXNS>
        <RESIDUE_SELECTORS>
        </RESIDUE_SELECTORS>
        <TASKOPERATIONS>
     </TASKOPERATIONS>
        <FILTERS>
        </FILTERS>
        <MOVERS>
          <SimpleGlycosylateMover name="glycans" glycosylation="a-D-Manp-(1->3)-[a-D-Manp-(1-
>3)-[a-D-Manp-(1->6)]-a-D-Manp-(1->6)]-b-D-Manp-(1->4)-b-D-GlcpNAc-(1->4)-b-D-GlcpNAc-"
positions="35,81,111,170,204,259,476,489,511,549,557,641,887,911,947,994,1041,1071,1123,1151,120
0,1212,1447,1460,1484,1522,1530,1614,1860,1884,1920,1967,2012,2039,2098,2131,2180,2192,2423,24
36,2457,2495,2503,2587,2833,2857,2893" idealize_glycosylation="true"/>
          <ParsedProtocol name="glyc_all">
            <Add mover_name="glycans"/>
          </ParsedProtocol>
        <GlycanTreeModeler name="glyc_relax" hybrid_protocol="1" shear="1"
use_gaussian_sampling="1" window_size="0" layer_size="1" match_window_one="1"
glycan_sampler_rounds="100"/>
        </MOVERS>
        <PROTOCOLS>
          <Add mover_name="glyc_all"/>
          <Add mover_name="glyc_relax"/>
        </PROTOCOLS>
</ROSETTASCRIPTS>
```
